# Quantifying dispersal between two colonies of northern elephant seals across 17 birth cohorts

**DOI:** 10.1101/2022.06.11.495743

**Authors:** Richard Condit, Brian Hatfield, Patricia A. Morris, Daniel P. Costa

**Affiliations:** Department of Ecology and Evolutionary Biology, University of California, Santa Cruz, USA; Institute for Marine Sciences, University of California, Santa Cruz, USA

**Keywords:** dispersal rate, elephant seal, recolonization, natal dispersal, pinniped

## Abstract

Dispersal drives extinction-recolonization dynamics of metapopulations and is necessary for endangered species to recolonize former ranges. Yet few studies quantify dispersal and even fewer examine consistency of dispersal over many years. The northern elephant seal (*Mirounga angustirostris*) provides an example of the importance of dispersal. It quickly recolonized its full range after near extirpation by 19^th^ century hunting, and though dispersal was observed it was not quantified. Here we enumerate lifetime dispersal events among females marked as pups at two colonies during 1994-2010, then correct for detection biases to estimate bidirectional dispersal rates. An average of 16% of females born at the Piedras Blancas colony dispersed northward 200 km to breed at Año Nuevo, while 8.0% of those born at Año Nuevo dispersed southward to Piedras Blancas. The northward rate fluctuated considerably but was higher than southward in 15 of 17 cohorts. The population at Piedras Blancas expanded 15-fold during the study, while Año Nuevo’s declined slightly, but the expectation that seals would emigrate away from high density colonies was not supported. During the 1990s, dispersal was higher away from the small colony toward the large. Moreover, cohorts born later at Piedras Blancas, when the colony had grown, dispersed no more than early cohorts. Consistently high natal dispersal in northern elephant seals means the population must be considered a single large unit in terms of response to environmental change. High dispersal was fortuitous to the past recovery of the species, and continued dispersal means elephant seals will likely expand their range further.

## Introduction

Dispersal and immigration are vital statistics of populations. Besides affecting gene flow and evolution, dispersal regulates metapopulation dynamics, and by overcoming local extinctions it can be crucial in species recovery from population crashes [1–3]. The northern elephant seal offers a clear example. Following near extermination by 19^th^-century hunters, it readily dispersed to reoccupy its former range [4–6]. During 50 years of research in California, we observed dispersal and documented new colonies formed by immigrants [7], but quantifying dispersal has been difficult, as is often the case in large animals that move long distances [8, 9]. Without precise estimates of rates of movement, we do not know whether dispersal continues now that the range is reestablished, and we cannot compare the importance of dispersal in elephant seals to other species [10, 11]. Here we fill the gap by quantifying dispersal rate in two directions in 17 consecutive birth cohorts. In a review of emigration rates, only four studies included more than 17 years [12].

Seals and other colonial animals have an important advantage in measurements of dispersal because breeding locations are discrete, so dispersal is a binomial process, either happening or not, and there are limited locations that must be searched to identify migrants. These features contrast with many terrestrial birds that have been the subject of dispersal research, where every individual disperses some distance [1, 9, 13]. We have been applying lifetime marks to female elephant seals for 30 years at two major colonies in central California, Año Nuevo and Piedras Blancas, and recording sightings of those animals at both colonies every year, so we can now be precise about dispersal in both directions. What percent of females born at one colony moved to the other to breed? Is there asymmetry in direction, with females more likely to move northward or southward? Our observations include the early phase of expansion at the Piedras Blancas colony, and so we can ask whether emigration increased as the colony expanded. Meantime, since Año Nuevo was large throughout, we can ask whether dispersal was greater from the large colony to the small, addressing the hypothesis that colony size drives emigration [12].

## Materials and Methods

### Ethics statement

Seal observations were authorized under Marine Mammal Research Permits 347, 404, 486-1506, 486-1790, 684, 704, 705, 774-1437, 774-1714, 836, and 14097; National Marine Sanctuary Permits GFNMS/MBNMS/ CINMS-04-98, MULTI-2002-003, MULTI-2003-003, and MULTI-2008-003; and Marine Mammal Protection Act Permit 486. Access to park land was granted by the California Department of Parks and Recreation.

### Study sites and colonies

Our study sites at Point Piedras Blancas and Año Nuevo (35.7^*◦*^, 37.1^*◦*^ N in California) are the largest northern elephant seal colonies on the mainland, providing accessibility and large samples. Other large colonies are on islands and difficult to access (Fig. 1). At both sites, female elephant seals gather in large groups on flat sand beaches every winter and give birth to a single pup. Pups are weaned an average of 26 days after birth when mothers depart to forage [14], and weaned pups are easily approached and tagged on the beach before they go to sea [15]. The Año Nuevo colony has had pups every winter since 1961. It expanded rapidly until 1995, then from 1995-2010, annual pup production declined slowly from 2700 to 2100 [6]. The Piedras Blancas colony first had pups in 1992 then grew from 300 pups in 1994 to 4400 in 2010 [6].

**Figure 1.** Map showing colony locations. The two study colonies at Año Nuevo and Piedras Blancas are marked with filled red circles. Other nearby colonies are marked with small triangles. The closest colonies at Southeast Farallon and Gorda are small, with only *∼* 100 breeding females each; Point Reyes has *∼* 900; the Channel Islands are enormous, with *>* 20, 000 at San Miguel and Santa Rosa combined [6]. There are three more large colonies much further south and another small one further north [6]. Axes show degrees latitude and longitude.

### Tagging and lifetime breeding records

Plastic sheep tags were inserted in the interdigital webbing of the hind flippers of weaned pups [15]; since 1998, most tags deployed were Jumbo Roto tags. On average, 21% of weaned pups were tagged at Año Nuevo and 11% at Piedras Blancas (Supplemental Table S1). We consistently searched both colonies for tagged adults during the winter breeding season, when females with pups hold their ground and allow observers closely. Tags were generally read with binoculars 3-5 m away from animals at Año Nuevo and with telescopes from bluffs 5-10 m away at Piedras Blancas. For this study we focus on pups tagged during 1994-2010. Because females start breeding at age 3-4 [16, 17], observations from 1997-2018 provide multiple opportunities to observe those cohorts during their breeding lifetimes. We assumed any female age 3 or older observed during the winter was breeding because 97.5% of adult females in the colony give birth [17, 18].

### Natal dispersal rate

Our focus is natal dispersal of females, defined as movement from the birth colony to a different colony on the first breeding attempt. The natal dispersal rate is the probability that a female alive for a first breeding attempt is at a colony different from her birth place. Any movements after initial breeding are termed adult dispersal, treated separately.

To estimate natal dispersal, define *T*_*i*_ as the total number of females tagged at colony *i*, and *B*_*i*_ the number of those observed breeding at any time in the future (Table S2 lists mathematical symbols). Now consider *b*_*ij*_ as the subset of *B*_*i*_ first observed breeding at colony *j*. For breeding residents, ie not dispersing, *i* = *j*, while in dispersers *i* ≠ j. In this study, *i, j ∈* (1, 2), and *B*_1_ = *b*_11_ + *b*_12_, *B*_2_ = *b*_21_ + *b*_22_. The ratio

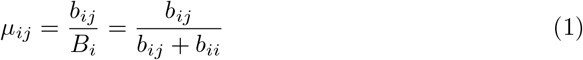

is a measure of natal dispersal rate from colony *i* to colony *j*. Any of the original *T*_*i*_ females never seen breeding do not enter into the calculation since we do not know whether or not they dispersed.

### Adult dispersal

Our goal is an analysis of natal dispersal, but we must address adult dispersal, defined as the movement between colonies after females begin breeding. If females move often as adults, estimates of natal dispersal will be confounded by adult dispersal in case females are not observed in their first breeding year. Preliminary calculations, though, showed adult dispersal to be rare [19], meaning that natal dispersal is effectively permanent dispersal: females spend entire lifetimes at a single colony. To confirm this, we present here an expanded estimate of adult dispersal based on observations of marked females across the 17 study cohorts. We found every case where a female was observed in consecutive breeding seasons and tallied the fraction of those that were at each of the two colonies. The proportion at a different colony in the second year is a direct estimate of annual adult dispersal.

### A detection bias

A concern with the calculation of dispersal is the failure to detect females that are present and breeding. Since our definition of natal dispersal is effectively lifetime emigration (given rare adult dispersal), it is lifetime detection that matters: the probability that a female who breeds one or more times during her lifetime is observed at least once. The concern here is that a difference in detection between our two study colonies will introduce a bias in dispersal estimates.

A simple example illustrates. Consider one cohort from colony 1 that ends up with 100 females breeding over their lifetimes, 90 resident where they were born and 10 dispersing permanently to a second colony, and a parallel cohort at colony 2 with 90 breeding as residents and 10 dispersing to colony 1. Both colonies have 10% dispersal. What if only half the females are detected over a lifetime at colony 1 but 80% at colony 2? Then the observed number of breeders born at site 2 and resident at site 2 would be *b*_22_ = 90×0.8 = 72, and the observed number emigrating would *b*_21_ = 10×0.5 = 5, leading to an estimate of dispersal from colony 2 to 1 of *µ*_21_ = *b*_21_*/*(*b*_21_+*b*_22_) = 5*/*(5+72) = 0.065. The opposite calculation leads to *µ*_12_ = *b*_12_*/*(*b*_12_ + *b*_11_) = 8*/*(8 + 45) = 0.15. Dispersal toward the colony with fewer observations will be underestimated, and vice versa.

We can correct for this bias if we know lifetime detection probability at each study site. At Año Nuevo, we know that females are not always detected, and annual detection – the probability that a tagged female alive during one winter season is observed – was *δ* = 0.6 in a mark recapture analysis [20]. Here we show that *δ* can be estimated from observations of marked females, thus providing separate values for our two colonies. Then we show how to propagate from annual to lifetime detection probability, defined as ∆. Correcting for any difference in ∆ leads to an unbiased comparison of dispersal between the two colonies.

### Estimating detection probability

Sightings of tagged females can be used to estimate annual and thus lifetime detection probabilities. First define rates of annual survival, *σ*, and tag retention, *ρ*. (Here we suppress subscripts, though every parameter is colony-specific; Table S2). In our definition, death includes permanent emigration outside the study colonies and tag loss means all tags are lost. There are previous estimates of survival and tag loss [18, 20], however, we demonstrate that they are not needed separately. Survival and retention appear in the calculations only as a product *τ* = *δρ*. Because it is the annual rate at which tagged females are back and observable after a year, we call it the return rate. Next, define a rate of reappearance, *π* = *δτ*, as the probability that a female returns with her tags and is detected. Thus

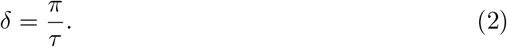

Both *τ* and *π* can be derived directly from observations. Appendices S1 and S2 give detailed derivations, but an intuitive grasp is straightforward. Return *τ* is the effective survival *σρ* (present and tagged), so it is the ratio of the number of animals alive at age *a* + 1 relative to a year earlier at age *a*:

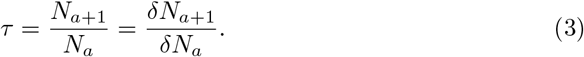

We cannot know either *N* directly, but we know *δN* because it is the number observed. Given the assumption that detection is constant from year to year and at all adult ages, we thus know the ratio *τ* . We overcome year-to-year variation in sighting effort by combining all ages across all years (Appendix S2).

The second required term, reappearance *π*, is the probability that an individual observed in one year is seen again the next year, because that requires survival, tag retention, and detection:

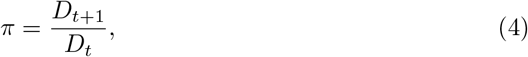

where *D*_*t*_ is the number of all animals observed in year *t* and *D*_*t*+1_ the subset of those also seen in year *t* + 1. The distinction between these two equations is that the numerator in Equation 4 includes animals seen in both years while in Equation 3 it includes all seen in the second year, even those not seen in the first. Thus detection emerges from the ratio.

Equation 2 produces annual detection, but we need lifetime detection ∆. Finding ∆ requires compounding multiple years of annual *δ*, ie the probability of failing to detect after two years is (1 − *δ*)^2^ etc. The full calculation (Appendix S3) leads to the following relation:

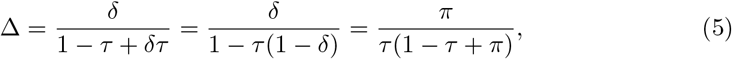

giving ∆ as a function of quantities known separately at each colony. The middle of the three forms is intended to offer some intuitive understanding, because the term 1*/*[1 − *τ* (1 − *δ*)] is the expected time until detection, showing that lifetime detection is annual detection multiplied by expected detection age.

### Correcting for lifetime detection

The number of tagged females observed breeding at least once over a lifetime can then be corrected for detection with

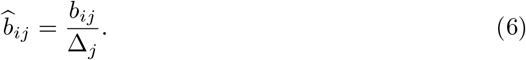

Since the subscript *j* identifies the colony where these animals were observed, the formula includes ∆_*j*_, lifetime detection at *j*. The corrected *b* leads to a correction for dispersal rate

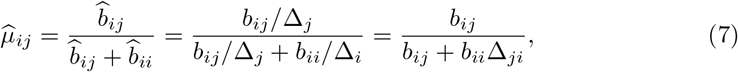

using Equation 1 and defining ∆_*ji*_ = ∆_*j*_*/*∆_*i*_ as the ratio of the detection probabilities. This shows that the correction term for dispersal is based solely on the ratio of the two detection probabilities, and that dispersal in the opposite direction depends on the inverse ∆_*ij*_ = 1*/*∆_*ji*_. Equation 7 is a quantitative statement of the qualitative conclusion that dispersal toward the well-observed colony is overestimated.

### Modeling and error propagation

A single estimate of the corrected annual dispersal is straightforward given Equations 3, 4, 5, and 7. But a thorough estimate of error requires propagating through all intermediate calculations. A Bayesian approach allows this.

First, there is error in the number of observed females *b*_*ij*_. We assume these are binomial draws from the total number breeding, *B*_*i*_ = *b*_*ii*_ + *b*_*ij*_, so that *b* is a Poisson random variable

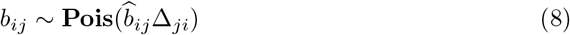

and thus has known error. The term in parentheses is the model’s prediction for the number observed given dispersal rate and detection (Eq. 6). In the Bayesian framework, Equation 8 is a likelihood function for observations (*b*) given a hypothesis (*b*_*ij*_∆_*ji*_, ie the model).

But there is also error in lifetime detection, and it is based on several intermediate calculations of the rates *π* and *τ* (Eq. 5). The first depends on a ratio of integer counts, so the sampling distribution is binomial and a posterior distribution of the ratio *D*_*t*+1_*/D*_*t*_ is beta,

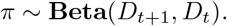

(The posterior is the probability of a given value of *π* given the data, the inverse of the likelihood function.) We created the posterior using the R function *rbeta* [21]. The second rate, *τ*, was derived from the slope of a regression from the age distribution (Appendix S1, Fig. S1). Standard linear regression leads to a slope *s* and its error *σ*_*s*_, thus

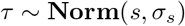

is a posterior probability of the desired parameter. We used the R function *rnorm* to generate it.

Here is where the value of the Bayesian approach arises. Random draws from the posteriors of *τ, π* were plugged into Equation 5 to produce a posterior distribution of lifetime detection at each colony, and those in turn were used to generate a posterior distribution of the ratio ∆_*ji*_. This final posterior became a prior probability in the model for dispersal (Eq. 7).

### A hierarchical model across years

A further feature we included with the Bayesian approach was to estimate a hyper-distribution across the annual dispersal parameters *µ*_*ij*_. This is a valuable tool here because individual years did not have large samples, and the hierarchy allows annual estimates to differ while still supporting each other [22]. We assumed the values *µ*_*ij*_ across years had a normal distribution described by hyper-parameters mean (*θ*_*ij*_) and standard deviation (σ_*ij*_),

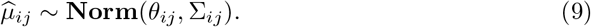

There are separate hyperparameters *θ*_*ji*_, σ_*ji*_ for dispersal in the opposite direction. We also tested a second multilevel model in which the mean *θ*_*ij*_ changed linearly through time, so *θ*_*ij*_ = *ν*_*ij*_ *· t* + *η*_*ij*_, where *t* is calendar year minus 2002. It has three hyper-parameters, slope *ν*_*ij*_, intercept *η*_*ij*_, and again σ_*ij*_ for the residual standard deviation. In the absence of a significant effect of year on dispersal (ie *ν*_*ij*_ not different from zero), we would prefer the simpler model with only *θ*_*ij*_, σ_*ij*_.

### Parameter estimation

The goal of the model is to generate estimates of dispersal parameters *µ*_*ij*_ for 17 cohorts, plus hyper-parameters *θ*_*ij*_, σ_*ij*_ (or *ν*_*ij*_, *η*_*ij*_, σ_*ij*_ for the regression version). Models for dispersal in two directions were separate, so two models each have 19 (20) parameters. Because there are several steps in calculations, and given the hierarchical aspect, generating posterior distributions for these parameters is not as easy as inverting a likelihood function. It required a Markov Chain Monte Carlo sampling method (MCMC) based on the Metropolis algorithm [23]. This meant repeated draws of all parameters, with the likelihood recalculated each time. The Metropolis algorithm is a tool for keeping the MCMC parameter draws in the vicinity of the maximum, producing precisely a posterior distribution for every parameter. MCMC is a standard Bayesian method [20, 24, 25].

Sampling the parameters requires an inverted version of Equation 7,

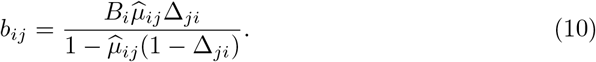

This gives the model prediction for the observed number of dispersers *b*_*ij*_ given the modeled *µ*_*ij*_ and ∆_*ji*_; *B*_*i*_ is known. Define **Θ** as the full vector of 19 (or 20) parameters per colony. The MCMC chain was started with a full set **Θ** using observed ratios *µ* for *µ*. At every step, each of the 17 annual *µ*_*ij*_ was plugged into Equation 10 (one-at-a-time) to predict 17 different *b*_*ij*_, the likelihood of each one found with Equation 8, then the likelihood of the hyper-parameters (given all 17 *µ*_*ij*_) calculated from Equation 9. Combining all likelihoods was accomplished by summing their logarithms. Each step of the sampler involved a new selection of all parameters **Θ** at random, one at a time, recalculating all likelihoods, then using Metropolis [23] to decide whether to adopt the new values or keep the previous.

Embedded within the sampler was the prior distribution for lifetime detection, ∆_*ji*_, already completed and stored. At each step of the MCMC, one value from this prior was drawn at random. Thus ∆_*ji*_ was updated along with the other parameters **Θ**, but did not depend on the observations *b*, only the prior. No prior probability was used on any of the other parameters **Θ**, except for the trivial restrictions that probabilities *µ ∈* (0, 1) and standard deviations σ *>* 0.

We executed samplers for 6000 steps and discarded the first 2000 as burn-in. Parameter chains converged quickly and consistently. The mean of post-burn-in chains was used as the best estimate for every parameter and the central 95^th^ percentiles as credible intervals.

### Inference on dispersal rate and colony size

The Piedras Blancas colony grew by 15-fold over the course of the study (Table S1). In early years, breeding females would have found relatively unoccupied beaches, in contrast to later years, when arriving animals found many beaches occupied. We do not know, however, whether colony density (females per beach area) increased with colony size, because animals spread out as the colony expanded. In contrast, the Año Nuevo colony was relatively stable, declining by 22% over the same period (Table S1), and seals occupied the same beaches throughout. We tested two hypotheses about the importance of colony size. First, we asked whether dispersal from Piedras Blancas increased through time using the hyper-parameter *ν*, the slope of the regression between dispersal and year. If its credible intervals overlapped zero, we would reject the influence of colony size. Second, we compared dispersal from Año Nuevo toward Piedras Blancas and vice versa using credible intervals to test the prediction that dispersal was more frequent from the large colony to the small. The hyper-means *θ* offered a test of the overall rates, plus there was a test every year based on credible intervals of the annual *µ*.

## Results

### Observed natal dispersal

Observed dispersal was more frequent from Piedras Blancas to Año Nuevo than vice versa (Table 1). Combining all 17 cohorts, there were 468 pups tagged at Piedras Blancas that were observed as breeding adults: 344 at their birth colony and 124 that emigrated to Año Nuevo (26.5% emigrated). There were 601 pups from Año Nuevo that were observed breeding: 566 as residents and 35 as emigrants to Piedras Blancas (5.8% emigrated). But this comparison is biased by detection, and the correction requires estimates of lifetime detection.

**Table 1.** **Observed dispersal within annual cohorts of female elephant seals. Tagged: total number of females tagged as pups each year at the two colonies. Breeding: the number of those observed breeding at least once over their lifetimes. Resident: subset of breeders that were first seen breeding where they were born. Emigrant: subset of breeders first seen breeding at the opposite colony.**

| Birth<br>year | Año Nuevo born |  |  |  | Piedras Blancas born |  |  |  |
| --- | --- | --- | --- | --- | --- | --- | --- | --- |
|  | Tagged | Breeding | Resident | Emigrant | Tagged | Breeding | Resident | Emigrant |
| 1994 | 240 | 24 | 24 | 0 | 120 | 13 | 11 | 2 |
| 1995 | 421 | 55 | 52 | 3 | 152 | 40 | 23 | 17 |
| 1996 | 271 | 51 | 46 | 5 | 148 | 41 | 33 | 8 |
| 1997 | 210 | 16 | 15 | 1 | 14 | 3 | 3 | 0 |
| 1998 | 211 | 37 | 36 | 1 | 155 | 35 | 23 | 12 |
| 1999 | 146 | 21 | 21 | 0 | 158 | 24 | 16 | 8 |
| 2000 | 264 | 46 | 45 | 1 | 156 | 32 | 14 | 18 |
| 2001 | 108 | 16 | 14 | 2 | 158 | 45 | 29 | 16 |
| 2002 | 138 | 40 | 39 | 1 | 158 | 42 | 36 | 6 |
| 2003 | 177 | 46 | 39 | 7 | 138 | 29 | 22 | 7 |
| 2004 | 168 | 25 | 23 | 2 | 150 | 31 | 31 | 0 |
| 2005 | 378 | 42 | 36 | 6 | 176 | 16 | 15 | 1 |
| 2006 | 286 | 28 | 28 | 0 | 132 | 20 | 15 | 5 |
| 2007 | 366 | 56 | 54 | 2 | 176 | 31 | 25 | 6 |
| 2008 | 214 | 26 | 25 | 1 | 174 | 22 | 18 | 4 |
| 2009 | 298 | 46 | 44 | 2 | 123 | 16 | 10 | 6 |
| 2010 | 208 | 26 | 25 | 1 | 196 | 28 | 20 | 8 |
| Total | 4104 | 601 | 566 | 35 | 2486 | 468 | 344 | 124 |

### Lifetime detection

At Piedras Blancas, the estimated lifetime detection was ∆ = 0.549, while at Año Nuevo, ∆ = 0.923, a highly significant difference (credible intervals in Table 2). The ratio of 0.598 (Table 2) means that the average female spending a lifetime breeding at Piedras Blancas was 60% as likely to be detected as a similar female at Año Nuevo. The observed reappearance rate, *π*, drove the difference: 16% of females seen in one year at Piedras Blancas were detected the next year, compared to 57% at Año Nuevo (Table 3). Since return rates *τ* – survival and tag retention – were indistinguishable between the colonies, the difference in annual detection matched the difference in reappearance (Table 2).

**Table 2.** **Parameter estimates from the Bayesian dispersal model. The first five parameters are colony-specific, but detection ratio and mean dispersal relate the two colonies and include subscripts 12 to show this. Under the Año Nuevo column, detection ratio is Año Nuevo divided by Piedras Blancas, while dispersal is from Año Nuevo to Piedras Blancas. The Piedras Blancas column has the opposite. For each, the best estimate is followed by 95% credible intervals in parentheses.**

| Rate | Symbol | Colony |  |
| --- | --- | --- | --- |
|  |  | Año Nuevo | Piedras Blancas |
| Reappearance | $\pi$ | 0.563 (0.54,0.58) | 0.165 (0.14,0.19) |
| Return | $\tau$ | 0.799 (0.79,0.81) | 0.783 (0.75,0.81) |
| Annual detection | $\delta$ | 0.704 (0.68,0.73) | 0.211 (0.18,0.25) |
| Lifetime detection | $\Delta$ | 0.922 (0.91,0.93) | 0.551 (0.49,0.61) |
| Year effect | $\nu$ | 0.012 (-0.09,0.11) | -0.024 (-0.11,0.06) |
| Detection ratio | $\Delta_{12}$ | 1.678 (1.52,1.87) | 0.598 (0.54,0.66) |
| Mean dispersal | $\theta_{12}$ | 0.0799 (0.05,0.11) | 0.1590 (0.11,0.22) |

**Table 3.** **Reappearance rate of breeding females. The total sample is the sum of all lifetime breeding records of females in the study cohorts through year 2017 (ie excluding 2018); any female seen more than once counts every time. The two rows divide the total sample based upon breeding location (birth location is not relevant here). For example, the 2297 breeding records at Año Nuevo is the sum of all records of 601 females born at Año Nuevo and 344 females born at Piedras Blancas (numbers in Table 1). There are a few individual females counted in both rows because of adult dispersal. Location in year 2 subdivides each row based on where females were observed one year later, none meaning not seen anywhere; they add up to the total. The reappearance rate, *π*, is the fraction reappearing at the same colony.**

| Location in<br>year 1 | Total<br>sample | Location in year 2 |  |  |
| --- | --- | --- | --- | --- |
|  |  | Año Nuevo | Piedras Blancas | None |
| Año Nuevo | 2297 | 1293 | 8 | 996 |
| Piedras Blancas | 671 | 2 | 110 | 559 |

### Adult dispersal

Of more than 1400 observations of breeding attempts in consecutive years, only 10 (0.7%) included females moving to the opposite colony in the second year (Table 3). Two moved from Piedras Blancas to Año Nuevo and eight the opposite.

### Corrected natal dispersal

Higher lifetime detection probability at Año Nuevo means that the observed number dispersing toward Año Nuevo (Table 1) is biased upward. Yet even after the correction, dispersal northward from Piedras Blancas was double that southward from Año Nuevo, 0.16 versus 0.08 (Table 2). The 95% credible intervals just met (Table 2).

There was considerable year-to-year variation in the rate from Piedras Blancas, indeed the rate of 0.41 for the cohort born in 2000 was significantly higher than the rate of 0.17 in 2004 (Fig. 2). There was much less variation in the southward direction, and none was significant (Fig. 2).

**Figure 2.** Natal dispersal by birth cohort. Dispersal rate of juveniles born at Piedras Blancas (PB) to breed at A∼no Nuevo (AN) in red. Dispersal from A∼no Nuevo to Piedras Blancas in blue. Best estimates are filled circles, and 95% credible intervals are vertical lines. The overall mean across all cohorts are the rightmost points with dashed credible intervals (Table 2).

### Dispersal and density

There was no temporal trend in dispersal in either direction (parameter *ν* did not differ from zero, Table 2). During the early years of the study, when the Año Nuevo colony had far more animals than Piedras Blancas, dispersal was higher from Piedras Blancas toward Año Nuevo, and the northward rate in 2000 was significantly higher than the southward (Fig. 2).

## Discussion

During the 1970s and 1980s, young elephant seals in California were observed emigrating from their birth colonies to establish new colonies [26, 27], but rates of dispersal were not estimated. Our results here confirm dispersal and add quantitative results over many years. Juvenile females dispersed between two major colonies 200 km apart at a substantial rate, 16% in the northward direction and 8% southward. Other published rates for pinnipeds are lower. In harbor seals (*Phoca vitulina*), 10% of breeding females emigrated to colonies within 10 km of their birthplace, but none dispersed more than 20 km [28], while in gray seals (*Halichoerus grypus*), the fraction dispersing was *<* 5% among all colony pairs, though that was based on population models, not direct observations [29]. Interestingly, a literature review covering many species (but no marine mammals), the mean fraction dispersing was 15% [10], close to what we observed. How dispersal was measured, and observed distances, were extremely variable among species, so this is not a safe generalization.

Dispersal rates between Año Nuevo and Piedras Blancas remained consistently high over 17 consecutive cohorts, and though there were fluctuations, there was no long-term trend. During that period, the population at Piedras Blancas grew 15-fold while the number at Año Nuevo declined slightly. There was thus no indication that emigration increased with colony size. Moreover, during the mid-1990s, there was more dispersal from the smaller Piedras Blancas colony toward the larger at Año Nuevo. These results do not support earlier observations that colony size drove decisions of young females to emigrate to found new colonies [7]. Those observations were based on the Año Nuevo Island colony, where females are tightly packed in one large group and most individuals encounter high density of near neighbors. In contrast, the Piedras Blancas colony spread to new beaches as it grew, so it is not clear whether individuals faced higher density after 2010 than in the 1990s. In a 2005 review of studies examining emigration and population density, Matthysen [12] found a positive relation in some studies but not others. A recent study of falcons (*Falco sparverius*) found no correlation between population density and frequency of dispersal [9].

If our study had ended with the cohort of 2005, we might have concluded that emigration from Piedras Blancas was declining after the high rates of 1995 and 2000, opposing the expectation that emigrants escape high density. Perhaps small size of the colony in the 1990s made it less attractive because elephant seals prefer dense aggregations. After 2005, however, emigration from Piedras Blancas rose again, and it is possible that positive and negative effects of density were both at work, or, more likely in our view, dispersal decisions are based on factors other than colony size.

There was directionality in dispersal, with the northward rate double the southward. The difference was at the margin of statistical significance, and it should be further tested, but the northward flow fits the elephant seal migration. Females move northwestward from their breeding colonies in California toward distant foraging grounds [30, 31], thus animals born at Piedras Blancas pass Año Nuevo on the migration, but not vice versa. The feeding grounds are considerably farther than the distance between colonies, and from that perspective dispersal distances in elephant seals are not high. A tendency for long-distance migrants to disperse well compared to non-migrants has been demonstrated in birds [32].

Our dispersal estimates include thorough analysis of error, most importantly the bias caused by unequal detection probabilities. This bias is often mentioned in studies of bird dispersal but is difficult to quantify [1]. We were able to estimate lifetime detection from many years of resightings of long-lived females, and we demonstrated much higher detectability at Año Nuevo than Piedras Blancas. Better detection at Año Nuevo can be attributed to better beach access as well as student participation in the university-sponsored research.

These natal dispersal rates may be underestimates because we only included two colonies, and we have evidence of movement to additional colonies. In Condit et al. [20], we reported the lifetime fate of three cohorts of females branded at Año Nuevo during the 1980s, including emigrants to two adjacent colonies. We found that seven emigrated of the 37 that bred (18.9%); this was before the Piedras Blancas colony existed, and the emigrants were at the Farallon and Point Reyes colonies north of Año Nuevo. In an analysis of three cohorts [19], all four colonies were included. No correction for detection was incorporated, but the observed number of emigrants from Piedras Blancas to the other two colonies was nearly as high as the number moving to Año Nuevo, while the number dispersing from Año Nuevo to the other two was even higher than the number moving to Piedras Blancas. Moreover, there may be dispersal to the Channel Island colonies, 200 km south of Piedras Blancas, but observations at those large island colonies are much more difficult. Overall, we have reason to believe that total natal dispersal rates are even higher than those reported here, perhaps reaching 20%.

Once breeding, adult females move at much lower rates. In the cohorts we studied, fewer than 1% moved over consecutive years. Northern elephant seal colonies are thus well-mixed genetically in that juveniles move among them at substantial rates, but most adults spend their lives at a single colony. Since all the colonies, including Mexico, form a chain with gaps of no more than 500 km, this mixing undoubtedly includes the entire range. In terms of response to the environment or prey abundance [33], we conclude that the northern elephant seal should be treated as one large population.

Perhaps most important is the role of dispersal in recovery from population bottlenecks, and this is clear in northern elephant seals. Nearly extinct in 1892 [4], the species refilled its range within 70-80 years [6, 34]. We demonstrate here that dispersal continues now, even after the range has filled, and elephant seals are thus poised to expand their range further, as they recently have at King Range in northern California [35]. In other pinnipeds, poor dispersal may be a primary reason for slow recovery [11]. The fact that elephant seals disperse well appears to be a fortuitous trait in the face of the 19^th^ century decimation by hunters.

## Supporting Information

Three Appendices provide complete derivations of the estimates of annual and lifetime detection of breeding females. Table S1 gives the number of animals tagged relative to those born at each colony, and Table S2 is a full list of mathematical symbols used. Figure S1 shows the rate of decay of the number of tagged females versus age at both colonies, used in estimating return rate *τ* .

## Acknowledgments

We are grateful to B.J. Le Boeuf for organizing and supervising the tagging program at Año Nuevo for many years. We also thank many colleagues, students, and volunteers for years of tagging and tag sighting at Año Nuevo; Brent Stewart and Ron Jameson for guidance; and the Friends of the Elephant Seal volunteers, especially Phil Adams, for tag observation at Piedras Blancas.

## Appendix S1: Rate of return

The first step in computing lifetime detection probabilities at the two colonies is an estimate of what we define as the rate of return of breeding females from year to year. This is the probability that a female alive in year *t* is still alive, has not emigrated, and has a tag in year *t* + 1, so return rate *τ* = *σρ* (Table S2). Under reasonable assumptions, *τ* can be estimated from observations of tagged animals. The annual detection term *δ* is not needed in these calculations given the assumption that *δ* is constant from year-to-year and throughout the breeding lifetime, until senescence. Lifetime survival curves suggest that both *σ* and *δ* are constant, with senescence setting in at age 17 [1]. Female behavior on the breeding colony is consistent at all ages 5 and above, also supporting a constant detection term. We do not have independent information on how tag retention, *ρ*, changes with age; we return to this later.

Start with a population of *N*_0_ females breeding for the first time. Then *D*_0_ = *δN*_0_ is the number of those observed. The subscript refers to an age, defining age=0 as the first year each female breeds. For animals one year older, the number returning (alive and tagged) is *N*_1_ = *σρN*_0_ = *τ N*_0_. The number of those detected is *D*_1_ = *δN*_1_ = *δτ N*_0_. At subsequent ages,

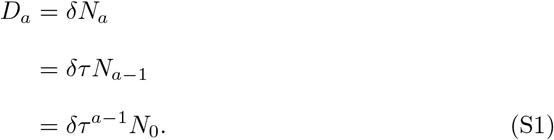

The exponent is *a* − 1 because we define the initial cohort as females alive and breeding for the first time at *a* = 0, so the survival probability until year *a* requires *a* − 1 survival steps. Taking logarithms,

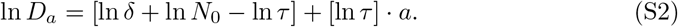

The two terms in square braces are constants that can be estimated from a regression of ln *D*_*a*_ against *a*. That is, the number of animals observed at successively greater ages declines exponentially with time, with slope ln *τ* (Fig. S1). We thus have a simple way of estimating *τ* . Detection *δ* appears in the intercept but not the slope based on the assumption that *δ* is constant during females’ breeding years, so its impact cancels out. Given the main breeding years described above, we calculated the age distribution from ages 5-15 to estimate this regression (Fig. S1). The regression slopes at the two colonies barely differed and were statistically indistinguishable (Table 2, main text). In both, slopes were slightly steeper after age 10, but not significantly so (Fig. S1). In Condit et al. [1], we demonstrated constant adult survival until age 17, and though we have not estimated tag loss in older animals, it evidently does not change much.

## Appendix S2: Annual detection

The next requirement for finding lifetime detection of breeding females is an estimate of annual detection probability, *δ*. This can be derived from observations of tagged females in consecutive years, given the estimate of return rate *τ* (Appendix S1). Here, define *N*_0_ as the number of tagged adult females breeding in year 0 (in contrast to Appendix S1, where the subscript referred to age, not year). We observe *D*_0_ = *δN*_0_ of those animals. A year later, the number of those *D*_0_ returning is *τ D*_0_, and using the assumption that detection probability is equal in the two years, the number of those detected is *D*_1_ = *δτ D*_0_ = *πD*_0_. The fraction *π* = *D*_1_*/D*_0_ is the reappearance rate as defined in the main text (Table S2). Then

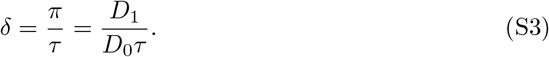

Notice that *D*_1_ is not all animals detected in year 1, it is only those detected in year 1 that had also been detected in year 0. It does not include animals observed in year 1 but not observed in year 0. In contrast, the successive ratios of Equation S2 and Figure S1 are based on all animals seen each year.

## Appendix S3: Lifetime detection

Now we derive the probability that a breeding female elephant seal is detected at least once in her lifetime, given that she bred at least once. Again, we use the annual return rate, *τ*, which is the product of annual survival *σ* and tag retention *ρ*, and annual detection *δ*. We rely on the assumption that all three are constant through the adult lifetime, until the age of senescence. We also must assume that a female’s detection in one year is independent of her detection in other years; we do not have evidence to support this assertion.

Female elephant seals can breed many years, and assuming independence from year to year, long-lived females will almost certainly be detected. Lifetime non-detection is most likely in females that breed once then die. The full derivation requires the probability of breeding once then dying, breeding twice then dying, etc.

It is easier to work with non-detection probabilities, so define annual non-detection as *λ* = 1 − *δ* and lifetime non-detection as Λ = 1 − ∆. The calculation begins with a group of females that are present on the breeding colony for the first time in their lives, *N*_0_, exactly as in Appendix S1. Consider the subset of this group that returns in exactly *n* breeding seasons, then dies (or emigrates or loses tags), so there are *n* chances to detect this group. Failure to detect in every one is *λ*^*n*^, so the probability of lifetime non-detection over exactly *n* years is

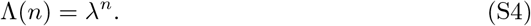

Since females have variable lifetimes, we need to calculate the probability of each lifetime *n*. Call *P* (*n*) the probability that the average female returns in exactly *n* breeding seasons, disappearing by *n* + 1. Because we assume the initial cohort was already alive in the first year, *n* − 1 return events are required. The probability is thus

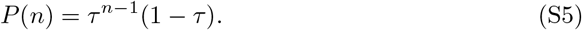

*τ* is the annual return rate (Appendix S1, Table S2).

We need this calculation for every lifespan: those females returning exactly *n* = 1, 2, 3, … years. Each of those groups includes a proportion of females given by Equation S5, and each is subdivided into two smaller groups, those not detected and those detected, the former from (Eq. S4). The product of Equations S4 and S5 is the proportion of females with a given lifetime that were never detected, so we sum those products over all lifetimes to find the total proportion never detected,

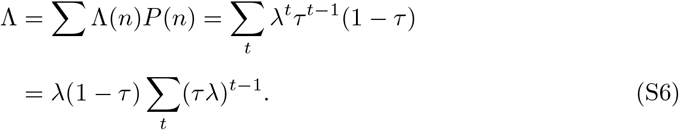

The summation runs from *t* = 1 to *t* = *∞* (years). The final version is rearranged so that the two terms inside the summation have the same exponent, *t* − 1. Then Equation S6 is an infinite geometric series, each term a factor *τλ* times the previous. In reality, the series would have to be curtailed when senescence starts, but the size of the seventeenth term is vanishingly small, so the infinite approximation is very close. Since we need lifetime detection, ∆ = 1 − Λ, the formula for a geometric series [2] yields

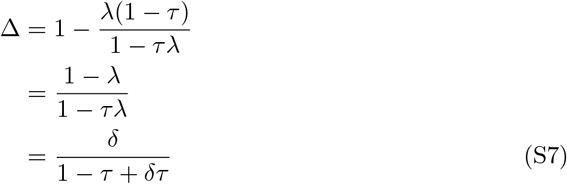

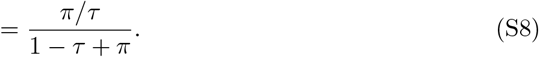

These give the lifetime detection probability as a function of the annual detection, *δ*, annual return, *τ*, and annual reappearance, *π*.

---

**Table S1.** **Number of females weaned and tagged each year of the study at both colonies, Año Nuevo (AN) and Piedras Blancas (PPB). Number weaned is from published tables [?, ?], divided by two (assuming females are half those weaned). The number tagged means females; it includes half of a small number whose sex was not recorded (Table 2, main text).**

| Year | Weaned |  | Tagged |  | Fraction tagged |  |
| --- | --- | --- | --- | --- | --- | --- |
|  | PPB | AN | PPB | AN | PPB | AN |
| 1994 | 146 | 1024 | 120 | 240 | 0.822 | 0.234 |
| 1995 | 302 | 1116 | 152 | 421 | 0.503 | 0.377 |
| 1996 | 494 | 1123 | 148 | 271 | 0.300 | 0.241 |
| 1997 | 598 | 1160 | 14 | 210 | 0.023 | 0.181 |
| 1998 | 836 | 1128 | 155 | 211 | 0.185 | 0.187 |
| 1999 | 956 | 1100 | 158 | 146 | 0.165 | 0.133 |
| 2000 | 923 | 1108 | 156 | 264 | 0.169 | 0.238 |
| 2001 | 970 | 1060 | 158 | 108 | 0.163 | 0.102 |
| 2002 | 1088 | 1124 | 158 | 138 | 0.145 | 0.123 |
| 2003 | 1324 | 1202 | 138 | 177 | 0.104 | 0.147 |
| 2004 | 1526 | 1010 | 150 | 168 | 0.098 | 0.166 |
| 2005 | 1757 | 1222 | 176 | 378 | 0.100 | 0.309 |
| 2006 | 1894 | 1201 | 132 | 286 | 0.070 | 0.238 |
| 2007 | 2040 | 1161 | 176 | 366 | 0.086 | 0.315 |
| 2008 | 2004 | 1072 | 174 | 214 | 0.087 | 0.200 |
| 2009 | 1904 | 1006 | 123 | 298 | 0.065 | 0.296 |
| 2010 | 2234 | 922 | 196 | 208 | 0.088 | 0.226 |

**Table S2.** **Math symbols used. Latin letters are observed or predicted counts of animals; Greek letters are parameters to be estimated (*σ* and *ρ* are only estimated as a product). Subscript *i* refers to colony; parameters with *i* alone are colony-specific. Those with *ij* relate two colonies, *i* and *j***

| Symbol | Definition |
| --- | --- |
| $T_i$ | Number of animals tagged as pups |
| $B_i$ | Number of $T_i$ later observed breeding (at any colony) |
| $b_{ij}$ | Number of animals born at $i$ then first observed at $j$ ($i = j$ or $i \neq j$) |
| $D_{ai}$ | Number of animals detected breeding, age (or year) $a$ |
| $N_{ai}$ | Number of animals alive, age (or year) $a$ |
| $\sigma_i$ | Annual survival rate |
| $\rho_i$ | Annual tag retention rate |
| $\delta_i$ | Annual detection probability |
| $\tau_i$ | Annual return rate ($= \sigma_i \rho_i$) |
| $\pi_i$ | Annual reappearance rate ($= \delta_i \sigma_i \rho_i$) |
| $\Delta_i$ | Lifetime detection probability |
| $\Delta_{ij}$ | Lifetime detection ratio, colony $i$ to $j$ |
| $\lambda_i$ | Annual non-detection probability ($= 1 - \delta_i$) |
| $\Lambda_i$ | Lifetime non-detection probability ($= 1 - \Delta_i$) |
| $\mu_{ij}$ | Uncorrected dispersal rate from $i$ to $j$ ($= b_{ij} / [b_{ij} + b_{ii}]$) |
| $\hat{b}_{ij}$ | Number of animals expected given lifetime detection ($= b_{ij} / \Delta_j$) |
| $\hat{\mu}_{ij}$ | Annual dispersal rate corrected for lifetime detection ($= \hat{b}_{ij} / [\hat{b}_{ij} + \hat{b}_{ii}]$) |
| $\theta_i$ | Hyper-mean of corrected dispersal (overall mean across years) |
| $\Sigma_i$ | Hyper-standard-deviation of corrected dispersal |
| $\nu_i$ | Slope from regression between annual dispersal rate and year |
| $\eta_i$ | Intercept from same regression |
| $\Theta_i$ | Vector of all parameters in model (annual dispersal, hyper-parameters) |

**Figure S1.**
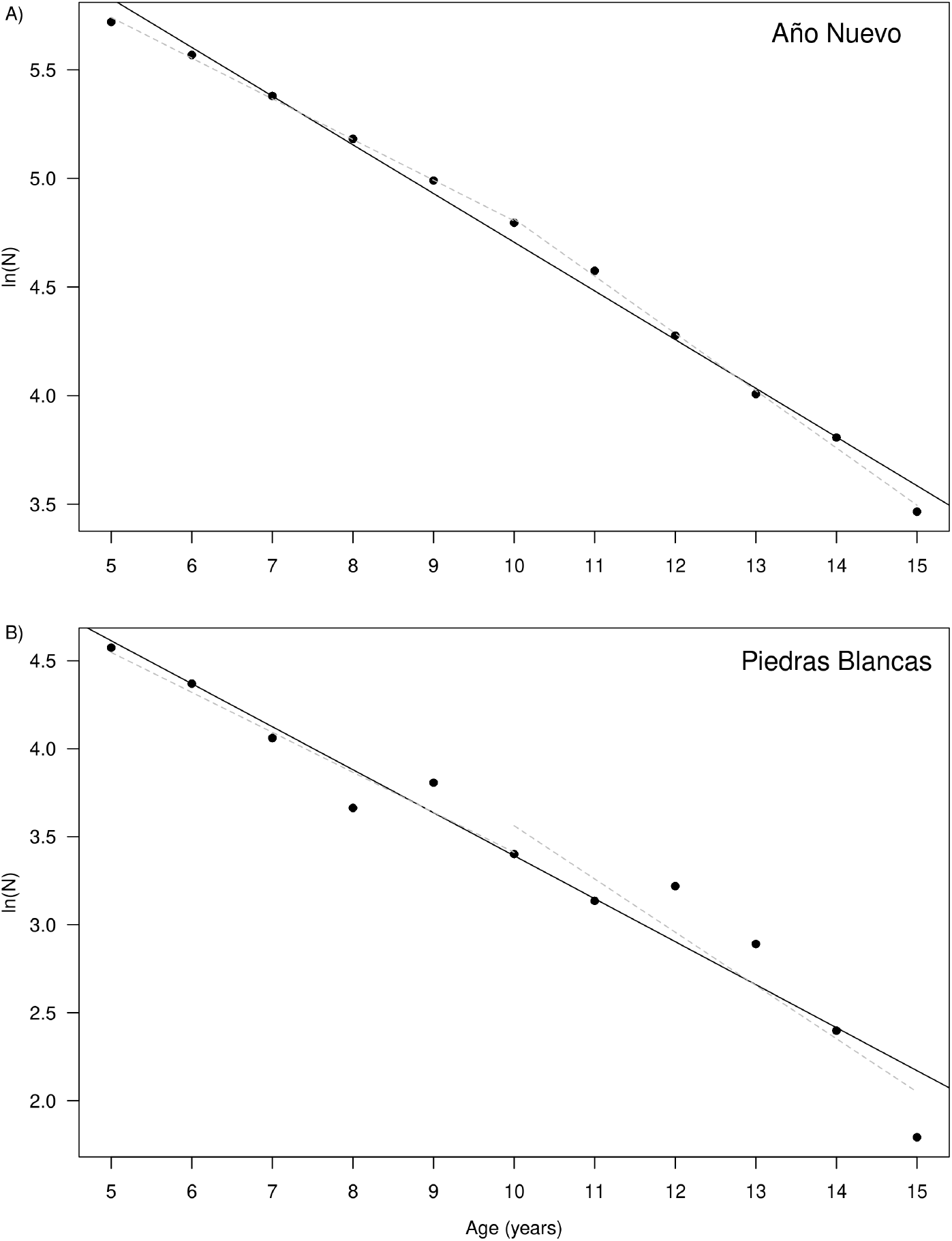
Decline in number of observed females on a colony with age. The vertical axis gives the natural log of the number of animals observed at each age, combining cohorts 1994-2010. The horizontal axis is female age in years; ages 5-15 include the main breeding life, when fecundity is at its maximum and before senescence. A) Año Nuevo. B) Piedras Blancas. The slopes provide estimates of ln *τ*, the logarithm of the rate of return (Appendix S1); *τ* = 0.799 at Año Nuevo (standard error 0.0056), *τ* = 0.783 (standard error 0.015) at Piedras Blancas. At each colony, a separate slope was fitted using ages 5-10 and again ages 10-15 (dashed gray lines). In both cases, the second slope was steeper, but in neither case did it differ statistically from the first.

## References

1. Winkler DW, Wrege PH, Allen PE, Kast TL, Senesac P, Wasson MF, et al. The natal dispersal of tree swallows in a continuous mainland environment. Journal of Animal Ecology. 2005;74:1080–1090.

2. Chilvers BL, Wilkinson IS. Philopatry and site fidelity of New Zealand sea lions (Phocarctos hookeri). Wildl Res. 2008;35(5):463–470.

3. Mestre A, Barfield M, Peniston JH, Peres-Neto PR, Mesquita-Joanes F, Holt RD. Disturbance-induced emigration: an overlooked mechanism that reduces metapopulation extinction risk. Ecology. 2021;102(8):e03423. doi:https://doi.org/10.1002/ecy.3423.

4. Townsend CH. The Northern Elephant Seal. New York Zoological Society; 1912.

5. Bartholomew GA, Hubbs CL. Winter population of pinnipeds about Guadalupe, San Benito, and Cedros Islands, Baja California. Journal of Mammalogy. 1952;33:160–171.

6. Lowry MS, Condit R, Hatfield B, Allen SG, Berger R, Laake J, et al. Abundance, distribution, and growth of the northern elephant seal, Mirounga angu-stirostris, in the United States in 2010. Aquatic Mammals. 2014;40(1):20–31. doi:10.1578/AM.40.1.2014.20.

7. Le Boeuf BJ, Condit R, Morris PA, Reiter J. The northern elephant seal rookery at Año Nuevo: a case study in colonization. Aquatic Mammals. 2011;37:486–501. doi:10.1578/AM.37.4.2011.486.

8. Tittler R, Villard MA, Fahrig L. How far do songbirds disperse? Ecography. 2009;32(6):1051–1061. doi:https://doi.org/10.1111/j.1600-0587.2009.05680.x.

9. McCaslin HM, Caughlin TT, Heath JA. Long-distance natal dispersal is relatively frequent and correlated with environmental factors in a widespread raptor. Journal of Animal Ecology. 2020;89(9):2077–2088. doi:https://doi.org/10.1111/1365-2656.13272.

10. Bowne DR, Bowers MA. Interpatch movements in spatially structured populations: a literature review. Landscape Ecology. 2004;19(1):1–20. doi:10.1023/B:LAND.0000018357.45262.b9.

11. Chilvers BL, Dobbins MLP. Behavioural plasticity and population connectivity: Contributors to the establishment of new pinniped breeding colonies. Aquatic Conservation: Marine and Freshwater Ecosystems. 2021;2021:1–12. doi:https://doi.org/10.1002/aqc.3603.

12. Matthysen E. Density-dependent dispersal in birds and mammals. Ecography. 2005;28(3):403–416. doi:https://doi.org/10.1111/j.0906-7590.2005.04073.x.

13. Greenwood PJ, Harvey PH. The Natal and Breeding Dispersal of Birds. Annual Review of Ecology and Systematics. 1982;13:1–21.

14. Reiter J. Studies of female competition and reproductive success in the northern elephant seal. University of California: Santa Cruz; 1984.

15. Le Boeuf BJ, Whiting RJ, Gantt RF. Perinatal behavior of northern elephant seal females and their young. Behaviour. 1972;43(1/4):121–156.

16. Reiter J, Le Boeuf BJ. Life history consequences of variation in age at primiparity in northern elephant seals. Behavioral Ecology and Sociobiology. 1991;28(3):153–160.

17. Le Boeuf B, Condit R, Reiter J. Lifetime reproductive success of northern elephant seals (Mirounga angustirostris). Canadian Journal of Zoology. 2019;97:1203–1217. doi:dx.doi.org/10.1139/cjz-2019-0104.

18. Le Boeuf BJ, Reiter J. Lifetime reproductive success in northern elephant seals. In: Clutton-Brock TH, editor. Reproductive Success. University of Chicago Press; 1988. p. 344–362.

19. Zeno R, Condit R, Allen SG, Duncan G. Natal and adult dispersal among four elephant seal colonies. BioRxiv. 2021;doi:10.1101/2021.03.18.435977.

20. Condit R, Reiter J, Morris PA, Berger R, Allen SG, Le Boeuf BJ. Lifetime survival and senescence of northern elephant seals, Mirounga angustirostris. Marine Mammal Science. 2014;30(1):122–138. doi:10.1111/mms.12025.

21. R Core Team. R: A Language and Environment for Statistical Computing; 2013. Available from: http://www.R-project.org/.

22. Gelman A, Hill J. Data Analysis Using Regression and Multilevel-Hierarchical Models. Cambridge University Press; 2007.

23. Metropolis N, Rosenbluth AW, Rosenbluth MN, Teller AH, Teller E. Equation of state calculations by fast computing machines. Journal of Chemical Physics. 1953;21(6):1087–1092.

24. Condit R, Ashton P, Bunyavejchewin S, Dattaraja HS, Davies S, Esufali S, et al. The importance of demographic niches to tree diversity. Science. 2006;313(5783):98–101. doi:https://doi.org/10.1126/science.1124712.

25. Condit R, Allen SG, Costa DP, Codde S, Goley PD, Le Boeuf BJ, et al. Estimating population size when individuals are asynchronous: a model illustrated with northern elephant seal breeding colonies. PLoS ONE. 2022;17(1):e0262214. doi:10.1371/journal.pone.0262214.

26. Le Boeuf BJ, Ainley DG, Lewis TJ. Elephant seals on the Farallones: population structure of an incipient breeding colony. Journal of Mammalogy. 1974;55:370–385. doi:https://doi.org/10.2307/1379005.

27. Allen SG, Peaslee SC, Huber HR. Colonization by northern elephant seals of the Point Reyes Peninsula, California. Marine Mammal Science. 1989;5:298–302.

28. Harkonen T, Harding KC. Spatial structure of harbour seal populations and the implications thereof. Canadian Journal of Zoology. 2001;79:2115–2127.

29. Harrison PJ, Buckland ST, Thomas L, Harris R, Pomeroy PP, Harwood J. Incorporating movement into models of grey seal population dynamics. Journal of Animal Ecology. 2006;75(3):634–645. doi:10.1111/j.1365-2656.2006.01084.x.

30. Aurioles D, Koch PL, Le Boeuf BJ. Differences in foraging location of Mexican and California elephant seals: evidence from stable isotopes in pups. Marine Mammal Science. 2006;22(2):326–338.

31. Robinson PW, Costa DP, Crocker DE, Gallo-Reynoso JP, Champagne CD, Fowler MA, et al. Foraging Behavior and Success of a Mesopelagic Predator in the Northeast Pacific Ocean: Insights from a Data-Rich Species, the Northern Elephant Seal. PLoS ONE. 2012;7(5):e36728.#x2013;.

32. Paradis E, Baillie SR, Sutherland WJ, Gregory RD. Patterns of natal and breeding dispersal in birds. Journal of Animal Ecology. 1998;67(4):518–536.

33. Goetsch C, Conners MG, Budge SM, Mitani Y, Walker WA, Bromaghin JF, et al. Energy-rich mesopelagic fishes revealed as a critical prey resource for a deep-diving predator using quantitative fatty acid signature analysis. Frontiers in Marine Science. 2018;5:1–19.

34. Stewart BS, Yochem PK, Le Boeuf BJ, Huber HR, DeLong RL, Jameson RJ, et al. Population recovery and status of the northern elephant seal, Mirounga angustirostris. In: Le Boeuf BJ, Laws RM, editors. Elephant Seals: Population Ecology, Behavior, and Physiology. University of California Press; 1994. p. 29–48.

35. Goley PD, Levy E. Northern elephant seal (Mirounga angustirostris) monitoring in the King Range National Conservation Area. Bureau of Land Management; 2021.

36. Purcell EJ. Calculus with Analytic Geometry; 1972.

## References

1. Condit R, Reiter J, Morris PA, Berger R, Allen SG, Le Boeuf BJ. Lifetime survival and senescence of northern elephant seals, Mirounga angustirostris. Marine Mammal Science. 2014;30(1):122–138. doi:10.1111/mms.12025.

2. Purcell EJ. Calculus with Analytic Geometry; 1972.

